# Canonical and noncanonical features of the mouse visual cortical hierarchy

**DOI:** 10.1101/2020.03.30.016303

**Authors:** Rinaldo D. D’Souza, Quanxin Wang, Weiqing Ji, Andrew M. Meier, Henry Kennedy, Kenneth Knoblauch, Andreas Burkhalter

## Abstract

Neocortical circuit computations underlying active vision are performed by a distributed network of reciprocally connected, functionally specialized areas. Mouse visual cortex is a dense, hierarchically organized network, comprising subnetworks that form preferentially interconnected processing streams. To determine the detailed layout of the mouse visual hierarchy, laminar patterns formed by interareal axonal projections, originating in each of ten visual areas were analyzed. Reciprocally connected pairs of areas, and shared targets of pairs of source areas, exhibited structural features consistent with a hierarchical organization. Beta regression analyses, which estimated a continuous measure of hierarchical distance, indicated that the network comprises multiple hierarchies embedded within overlapping processing levels. Single unit recordings showed that within each processing stream, receptive field sizes typically increased with increasing hierarchical level; however, ventral stream areas showed overall larger receptive field diameters. Together, the results reveal canonical and noncanonical hierarchical network motifs in mouse visual cortex.

## INTRODUCTION

The perception and interpretation of visual input is accomplished in part by generative intracortical and thalamocortical mechanisms that serve to provide contextual influences, focus attention, and compare incoming information with prior knowledge^1,2^. The classical notion of a cortical hierarchy posits that signals from primary visual cortex (V1) are routed through increasingly specialized cortical areas^3,4^. In such a hierarchical framework, ascending or feedforward pathways carrying sensory information are responsible for the retinotopic mapping of stimulus-selective receptive fields, whereas descending or feedback pathways selectively shape cortical responses within the receptive field depending on the context of the animal’s task or behavior^1,5,6^. In predictive coding theories of cognition, feedback pathways continuously convey predictions or estimates of a visual scene based on spatial and temporal statistical regularities within the scene, while incoming retinal signals are compared to these predictions; the residual error then ascends up the hierarchy for further comparisons with feedback predictions^7,8^. Accordingly, sensory representations are thought to be distributed across all of the functionally specialized areas that form the network, which has led to the view that visual perception is not a result of a chronological stepwise process in which visual signals are processed one area at a time^9-11^. It is unclear, therefore, if a sequential ranking of areas, as suggested by several hierarchical models^4,12-14^, fully describes the visual cortical network or whether the visual system utilizes noncanonical network strategies that involve communication between areas that do not conform to conventional feedforward and feedback relationships^15^. The term *noncanonical* is used here to describe an interconnected network in which reciprocally connected areas often violate feedforward/feedback relationships^15^, and such networks are therefore distinct from a strict sensory-motor hierarchy.

Because cortical lamination constrains circuit function^16^, across species the cortical hierarchy is determined by the laminar organization of interareal pathways. In primates, the termination of axonal projections in the target area, as well as the location of the cell bodies of the projecting neurons, show stereotypic laminar patterns depending on whether the projections are feedforward or feedback^4,17,18^. The distinct anatomical signatures of feedforward and feedback pathways have consequently been used to construct cortical hierarchies to identify a sequence of areas for the selective integration, amplification, and routing of visual signals^4,14,18-22^. Assigning a graded set of hierarchical level values to areas within a dense network^19,20^, as opposed to ranking areas based on pair-wise hierarchical relationships^4^, eliminates the need to classify individual projection patterns into feedforward or feedback categories. This is especially beneficial because the presence of *lateral* connections (i.e. connections that cannot be identified as either feedforward or feedback), violations of anatomical feedforward/feedback rules in reciprocally interconnected pairs, and missing interareal connections can all lead to multiple solutions for ordering areas. Indeed, using hierarchical level and distance measures, derived from anatomical criteria, has proven particularly useful in investigating the organization of primate cortex in order to solve the indeterminacy of the Felleman and Van Essen hierarchical diagram in which over 150,000 equally tenable solutions were estimated to exist^20,23^.

An exhaustive recent study, which analyzed the laminar termination patterns of axonal projections from 43 areas throughout the mouse neocortex, generated a hierarchy of the cortical network^14^. By designating a feedforward or feedback label to each laminar pattern such that the consistency of hierarchical relationships between areas is maximized, and by calculating a hierarchical score for each area, the analysis did not allow for connections to be categorized as being lateral, resulting in the construction of a linearly ordered hierarchy of areas^14^. However, given the ‘ultra-dense’ cortical graph of the mouse visual system in which almost all possible connections between visual areas have been shown to exist^24,25^, it is unclear to what extent consistent distance rules and hierarchical relationships govern the entire network; for example, a reciprocally connected pair may exhibit laminar patterns in both directions that are categorized as feedforward, as has been reported to exist in the macaque^19,20^. Further, despite dense interconnectivity, interareal visual pathways have been shown to exhibit preferences in the weight of connections between specific areas, leading to the observation that, like primates, mouse visual cortex is organized into the ventral and dorsal processing streams^25^. How processing streams are integrated into a hierarchy of areas is not well understood.

In this study, we aimed to deduce whether areas involved in visual representation fit into a consistent, functional sequence of distinct processing levels that communicate with higher and lower levels through unambiguous feedforward and feedback pathways, or whether the network is noncanonically organized. We examined the laminar termination patterns of cortico-cortical projections between V1 and nine previously identified higher visual areas that each receives retinotopically organized projections from V1, and whose borders were identified in each animal by anatomical landmarks^26^. By employing a beta regression model to determine hierarchical *level* and *distance* values, which best predict laminar termination patterns for each interareal connection, we show that while mouse visual cortex exhibits the features of an overall hierarchy, the network is characterized by a large number of lateral connections. The analyses indicate that the visual cortical graph can be best described as comprising four levels, with the highest level itself an interconnected network of areas showing hierarchical characteristics within an overall noncanonical framework. We further show using single unit recordings that receptive field sizes typically increase with increasing levels, but show a dependency on whether the respective areas belong to the dorsal or ventral stream.

## METHODS

All experimental procedures were approved by the Institutional Animal Care and Use Committee at Washington University in St. Louis.

### Animals

6-16 weeks old C57BL/6J male and female mice were used for the analyses of interareal axonal projection patterns. The *in vivo* single unit recordings were performed in 5-8 weeks old C57BL/6J male and female mice.

### Tracing axonal projections

A single injection of anterograde tracer was performed in each animal. Each mouse was anesthetized using a mixture of ketamine and xylazine (86 mg.kg^-1^/13 mg.kg^-1^, IP), and thereafter secured in a head holder. To retrogradely label callosally projecting neurons for areal identification in the left hemisphere, 30-40 pressure injections (Picospritzer, Parker-Hannafin) of bisbenzimide (5% in water, 20 nl each; Sigma) were made into the right hemisphere (occipital, temporal, and parietal cortices) using glass pipettes (20-25 µm tip diameter). Interareal connections within the left hemisphere were anterogradely labeled by inserting a glass pipette (15-20 µm tip diameter) into one of ten cortical areas and performing iontophoretic injections (3 µA, 7 s on/off cycle for 7 minutes; Midgard current source; Stoelting) of biotinylated dextran amine (BDA; 10,000 molecular weight, 5% in water, Invitrogen). Injections of BDA were performed stereotaxically and at two depths (0.3 mm and 0.5 mm) below the pial surface. The stereotaxic coordinate system had its origin at the intersection of the midline and a perpendicular line drawn from the anterior-most part of the transverse sinus at the posterior pole of the occipital cortex. The coordinates of the injected areas were (anterior/lateral in mm): V1, 1.1/2.8; LM, 1.4/4.0; AL, 2.4/3.7; RL, 2.8/3.3; PM, 1.9/1.6; P, 1.0/4.2; A, 3.4/2.4; LI, 1.45/4.2; AM, 3.0/1.7; POR, 1.15/4,3. Post-surgery analgesia was provided by subcutaneous injections of buprenorphine (0.05 mg.kg^-1^).

### Histology

At least 3 days after BDA injections, the mice were overdosed with ketamine/xylazine (500 mg.kg^-1^/50 mg.kg^-1^, IP) and perfused through the heart with heparin (0.01% in PBS, pH 7.4), followed by perfusion of 4% paraformaldehyde (PFA, in 0.1 M phosphate buffer [PB]), and brains were thereafter equilibrated in sucrose (30% in 0.1 M PB). To confirm the location of the injection site and to enable the identification of areal targets of axonal projections, bisbenzimide labeled neuronal landmarks in the left hemisphere were imaged *in situ* under a stereomicroscope (Leica MZ16F) equipped for UV fluorescence (excitation/barrier 360 nm/420 nm). The left hemisphere was then cut in the coronal plane on a freezing microtome at 40 µm. Sections were recorded as a complete series across the caudo-rostral extent of the hemisphere, and each wet-mounted section (in 0.1 M PB) was imaged under UV illumination using a fluorescence microscope equipped with a CCD camera (Photometrics CoolSNAP EZ, Sony). BDA labeled fibers were visualized by incubating the sections in avidin and biotinylated HRP (Vectastain ABC Elite), enzymatically reacted in the presence of diaminobenzidine (DAB) and H_2_O_2_, and the reaction product was intensified with AgNO_3_ and HAuCl_2_ ^25^. Sections were mounted onto glass slides, cleared with xylene, coverslipped in DPX, and imaged with dark field optics. Layers were assigned by the size and density of dark cell bodies on a brighter background, which resembled the patterns seen in contrast-inverted Nissl stained sections.

### In vivo electrophysiology

For single-unit recordings to measure receptive field (RF) sizes, mice were first anesthetized with urethane (20%, 0.2 ml/20 g body weight, i.p.). Tungsten electrodes dipped in DiI (5% in absolute ethanol) were inserted into each of the ten visual areas, guided by stereotaxic coordinates, in the left hemisphere. Recording depth was measured from the pial surface, and electrode insertion was controlled and monitored using a micromanipulator (Sutter Instruments). Recordings of spiking activity were acquired using the TEMPO software (Tempo, Reflective Computing), and single units were isolated through the use of a digital spike discriminator (FHC Inc.). Recorded signals were amplified and bandpass filtered at 300-5000 Hz.

RF location was first determined by moving a light bar over a dark background of the monitor screen, and listening to the audiomonitor response to spike discharges. To measure RF sizes, a circular patch (5 deg diameter) of a drifting sinusoidal grating (0.03 c/deg) was presented at different locations on the display screen. Spatial response plots were generated from contour lines connecting points in visual space with similar mean response strengths to visual stimuli. The response strength for a neuron was measured as the mean firing rate during the 2 s stimulus. For each recorded neuron, the response field was fit with a Gaussian, and the RF diameter was computed from the contour corresponding to two standard deviations (SDs) of the fitted Gaussian after transforming the elliptical field into a circle.

#### Confirmation of recording site

After each recording, mice were perfused through the heart with heparinized PB followed by 1% PFA (see ***Histology*** for details). The left cortical hemisphere was separated from the rest of the brain, flattened, postfixed in 4% PFA, and cryoprotected in 30% sucrose^26^. The flattened cortices were sectioned at 40 µm in the tangential plane on a freezing microtome. The sections were washed with 0.1 M PB, treated with 0.1% Triton X-100 and normal goat serum (10% NGS in PBS), and incubated with an antibody against the M2 muscarinic acetylcholine receptor (1:500 in PBS, MAB367, Millipore) for at least 24 hours. The sections were next washed with 0.1 M PB, and treated with a secondary antibody labeled with Alexa Fluor 647 (1:500 in PBS, Invitrogen). Sections mounted on glass slides were imaged with a CCD camera (Photometrics CoolSNAP EZ), and the location of the recording site marked with DiI was compared with the M2 staining pattern. The area location of the recording site was assessed using published maps based on M2 expression^27,28^.

### Data analysis and statistics

For each injected animal, 3-4 adjacent coronal sections containing anterogradely BDA labeled projections in each target area were used for quantification of termination patterns. Occasionally, the terminal projections contained retrogradely labeled cells. Such projections were excluded from analysis. Projections were assigned to areas by their location relative to bisbenzimide-labeled callosal landmarks^,21,26^ and based on their locations relative to other projection sites. BDA labeled projection fields were then superimposed over the pattern of bisbenzimide labeled callosal patterns in the same section, and axonal terminations were assigned to specific areas according to previously published maps^26^.

For analyses of the termination pattern in each area, each coronal section was imaged under 10x magnification, and grayscale images recorded. Contour plots of the optical density of BDA labeled axons were generated using a custom written MATLAB script^21^. Briefly, a circular averaging 2-D filter was used to blur the raw image, and contours denoting distinct optical density levels were generated. As explained in the Results section, layers (L) 1 to 4 were used for analyses. Regions bordered by the highest 70% optical density contour in L1 and L2-4 were used to measure the L2-4:L1 optical density ratio (ODR) for each slice. Optical densities were calculated using ImageJ.

#### Estimation of hierarchical levels

Statistical tests for generating and analyzing the cortical hierarchy were performed in the Open Source software R^29,30^. The regression analysis used here is an adaptation of the previously reported model that used discrete counts of retrogradely labeled cells instead of the continuous measure of optical densities of anterogradely labeled fibers used here for quantifying hierarchical relationships^20^.

We used beta regression to estimate hierarchical distance values between a pair of areas that would best predict the ODR for pathways connecting the two areas. The beta distribution is a continuous probability distribution defined on the interval (0, 1) typically parameterized by two shape parameters, *α* and *β*, with a probability density function

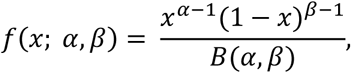

where 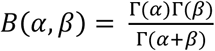, where Γ is the Gamma function. In the betareg package^29^, the distribution is reparameterized in terms of the mean, 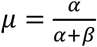, and a precision parameter, *ϕ* = *α* + *β*, with probability density

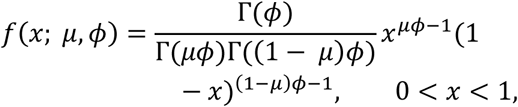

where 0 < *μ* < 1 and *ϕ* > 0.

To fit a hierarchical model, we estimate the expected value of the ODR and set the natural logarithm of the ODR equal to the hierarchical distance between two areas. This defines a relation between a linear predictor and the log of the ODR

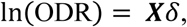

where ***X*** is the incidence matrix for the graph of areal connections, i.e., each column corresponds to one of the 10 areas and each row corresponds to a link between a pair of areas. Each row is composed of 0’s except for the columns corresponding to the connection between the two areas, taking the values - 1 and 1, depending on whether they are the source or target. The vector *δ* contains the estimated hierarchical levels assigned to each area (column). All of the rows sum to zero. Therefore, in order to yield an identifiable solution, one column is dropped - here the column corresponding to area V1 - and its hierarchical level is, thereby, fixed at 0. Provided the incidence matrix and the vector of ODR values, the betareg function returns maximum likelihood estimates of the vector *δ* and the precision parameter, *ϕ* and their standard errors estimated from the variance-covariance matrix. The solution is unique only up to addition of a constant and/or multiplication by a factor. The logit transformation can be inverted to yield the predicted proportion of axonal projections in L2-4 to the total axonal projections in L1-4

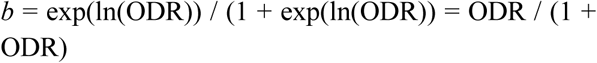

to compare with the empirical estimates. The inverse logit of *b* was provided by the *plogis* function in R, and the beta regression calculated using the *betareg* function in the betareg package^29^.

The Akaike Information Criterion (AIC)^31^ was used to assess the number of levels that best described the hierarchy. The AIC is defined as minus twice the log likelihood plus twice the number of parameters estimated. In the large sample limit, it approximates the same result as the leave-out-one cross-validation score, thus giving a measure related to prediction error. The model with the lowest AIC will yield the best balance of goodness of fit and model complexity and among a set of models is expected to best predict new data sets. The AIC values for the different models (see Results) were obtained using standard methods in R.

## RESULTS

### Diverse laminar projection patterns across cortico-cortical pathways

Feedforward and feedback axonal projections in the mouse visual cortex have been shown to exhibit distinct laminar termination patterns^14,21^. To observe these termination patterns for all connections within the network of visual areas, we injected the anterograde tracer BDA into V1 and each of nine higher visual areas that have been previously identified through topographic mapping of projections from V1^26^, and whose borders have been identified through anatomical and molecular landmarks^25,27^: LM (lateromedial), AL (anterolateral), RL (rostrolateral), P (posterior), LI (laterointermediate), PM (posteromedial), AM (anteromedial), A (anterior), and POR (postrhinal). One such injection was performed in each animal, and at least two animals were used for an injection in each area (n = 23 animals). To minimize BDA uptake by broken fibers of passage, and to ensure labeling of neurons in all six layers, iontophoretic injections with fine pipettes were performed through the depth of cortex. For each injection, laminar patterns of axonal projections in the nine target areas were examined in the coronal plane (Figure 1 a-c, Supplementary Figures 1 to 9). 40 µm coronal sections were numbered beginning from the posterior pole of occipital cortex, and individual areas were subsequently identified by their locations relative to landmarks formed by bisbenzimide labeled, callosally projecting neurons (see Methods; Supplementary Figure 1a) and by their locations relative to each other. An example injection in area AL is illustrated in Supplementary Figure 1. The callosal projection landmarks observed *in situ* were identified in coronal sections (Supplementary Figure 1a), which aided in the identification of areas. Similar images for injection sites *in situ* in areas V1, LM, and PM have been previously reported^21^. Representative termination patterns in target areas for injections in V1, LM, PM, RL, P, LI, A, AM, and POR are respectively shown in Figures 1a and Supplementary Figures 2 to 9. While all six layers were frequently targeted by interareal projections, the termination patterns exhibited striking laminar differences across pathways (Figure 1a-c, Supplementary Figures 1 to 9). Projections between higher visual areas often targeted all six layers, with varying relative density of fibers between layers. For example, Supplementary Figures 1a and 1b show that afferents from AL were densest in layer (L) 1 and the middle layers comprising L2/3 and L4 (L2-4) in A, AM, PM, and RL, while showing a preference for targeting L1 over all other layers in areas V1 and LM. AL also densely innervated L5 of areas A, AM, and RL.

**Figure 1.**
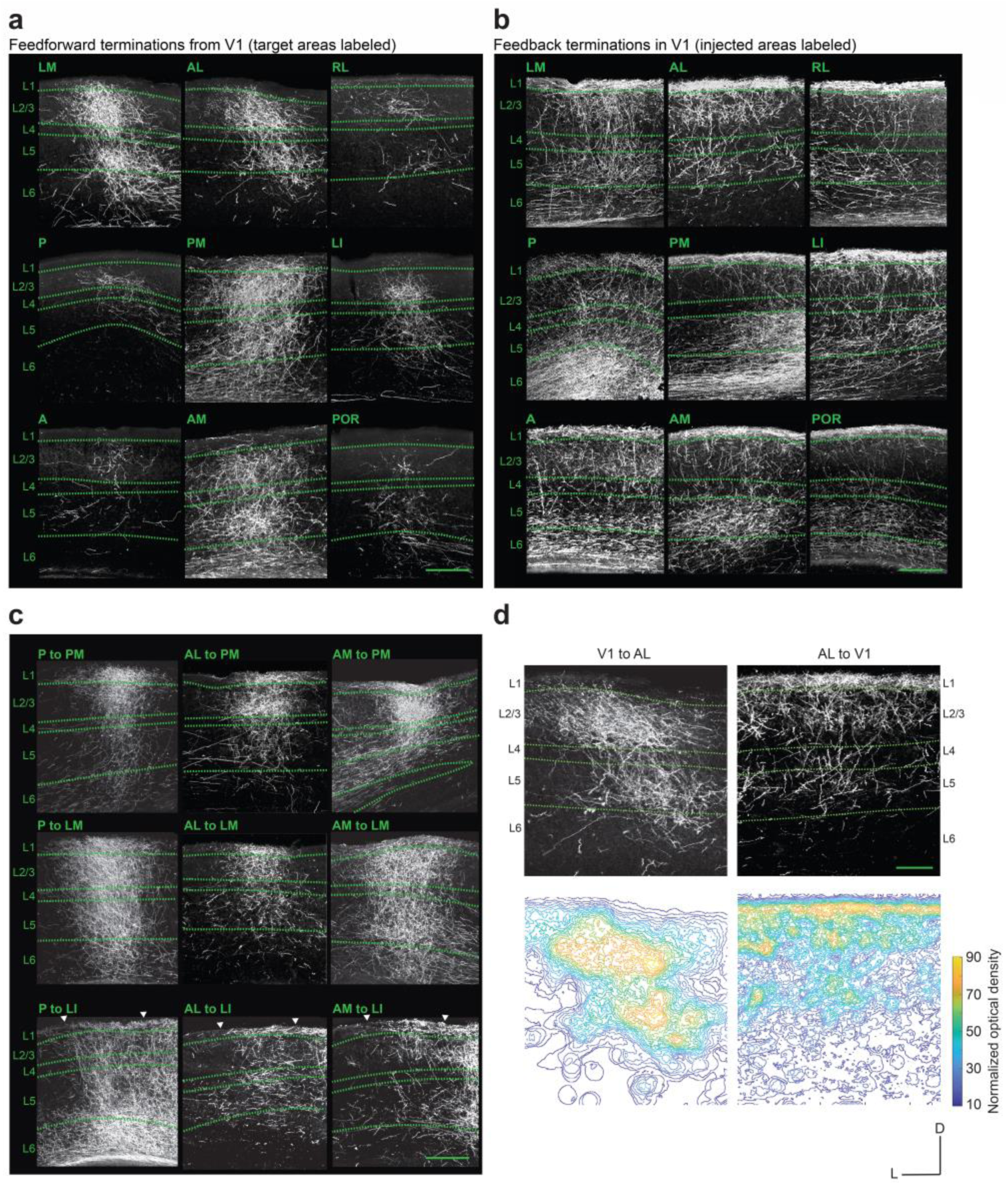
Darkfield images of coronal sections showing diverse laminar termination patterns of intracortical axonal projections. **a.** Laminar termination patterns of feedforward axonal projections in each higher-order area after injection of BDA into V1. **b.** Laminar termination patterns of feedback projections in V1 after injection of BDA into each of the nine higher-order areas. Injected areas are denoted for each image. One injection was performed in each animal. **c.** Nine representative examples of higher-order cortico-cortical laminar termination patterns, for injections of BDA performed in areas P, AL, and AM. Arrowheads demarcate boundaries of LI. Scale bars (a-c), 200 µm. **d.** Illustration of the generation of optical density contours for two example pathways. The average optical density in L1 and in L2-4 for each termination pattern was used for analysis. Scale bar, 100 µm.

To identify the respective anatomical signatures of feedforward and feedback pathways, we first focused on the termination patterns of projections emanating from, and terminating in, V1. As the primary geniculo-cortical target, V1 can be regarded as the lowest area of the visual cortical hierarchy; accordingly, projections originating in V1 and terminating in the other areas were classified as feedforward projections, whereas those targeting V1 and originating in the other areas as feedback. V1 projections terminated densely in regions encompassed by L2-4, with a relatively sparser termination in L1, of the target areas (Figure 1a). In contrast, feedback pathways to V1, originating in each of the higher areas, terminated most densely in L1, with substantially weaker projections to L2/3 and L4 (Figure 1b). Both feedforward and feedback pathways frequently exhibited dense terminations in L5 and L6; however, in addition, L6 often exhibited a high density of axons of passage traversing from the injection site to target areas (Figure 1a, b, Supplementary Figure 1a).

### Hierarchical characteristics in the mouse visual cortical network

Similar to previous analyses^21^, in order to quantify the polarity (i.e. whether feedforward or feedback) of pathways, we measured the optical density ratio (ODR), defined as the ratio of the optical density of axonal terminations in L2-4 to that within L1, for each connection. This ratio was chosen because the density of projecting axons in L2-4, relative to that in L1, provided a clear distinction of feedforward and feedback pathways, i.e. projections from and to V1, respectively. We therefore reasoned that the ODR would provide a graded hierarchical index that scales across pathways from the most feedforward to the most feedback. While both feedforward and feedback pathways commonly exhibited terminations in L5 and L6, these deep layers were often sites for axons of passage for interareal connections^32^, and stereotypic differences between feedforward and feedback connections to these layers could not be identified. Out of the 90 possible cortico-cortical connections between any two of the ten areas, 80 pathways exhibited projections in the target area, which were dense enough for analysis.

To calculate the ODR, optical density maps of BDA labeled axonal fibers in the target area were generated after running the image through a circular averaging filter (see Figure 1d for an example of density maps for the V1→AL and AL→V1 pathways), and the average optical density of regions within the highest 70% contours in L1 and L2-4 (i.e., the densest 70% of projections in these layers), after subtraction of background optical density, was computed. For each pathway, the ODRs from 3 to 5 coronal sections containing the center of the projection to the target area were averaged.

Upon analyzing these termination patterns, a 10 x 10 matrix of average ODRs for the 80 connections was generated (Figure 2a). Figure 2b shows the range of ODRs of laminar patterns from all coronal sections, and the asymmetric distribution of ODR values around 1. The frequency distribution of log ODR values exhibits a more symmetrical distribution (Figure 2b inset). Unsurprisingly, the matrix showed that feedforward projections from V1 had, on average, the largest ODR values, while feedback pathways to V1 exhibited the lowest (Figure 2a). Among the higher visual areas, projections from area LM had, on average, the highest ODR. This is consistent with the notion that LM may be considered the mouse analog of the secondary visual cortex (V2) of primate^25^.

**Figure 2.**
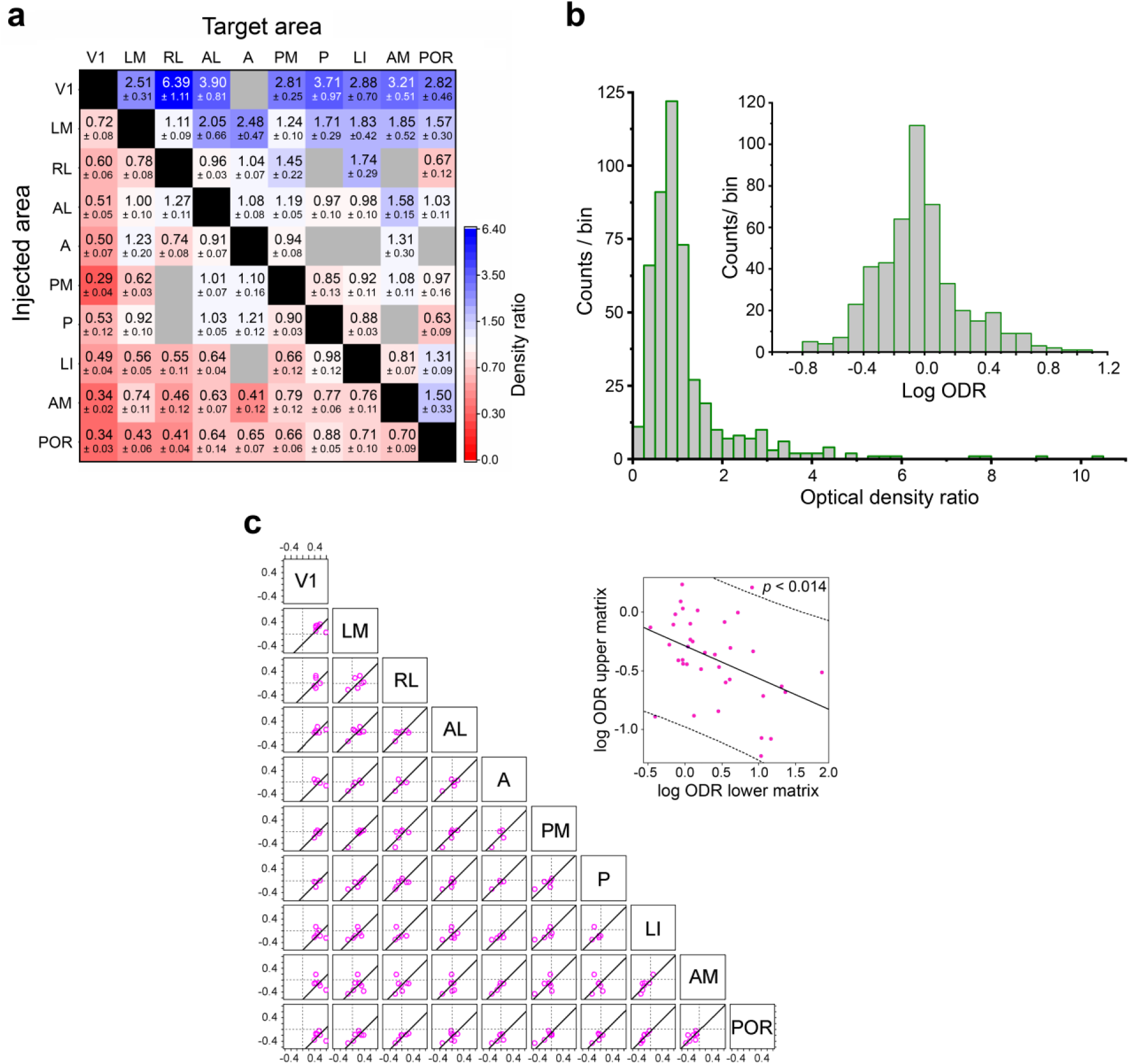
Mouse cortical network exhibits hierarchical features. **a.** 10 X 10 connection matrix of interareal connections between the ten visual areas. Each block shows the ratio of the optical density of axonal projections in L2-4 to that in L1 (optical density ratio, ODR) for the respective pathway in which the source and target areas are denoted on the vertical and horizontal axes, respectively. Grey blocks represent pathways that could not be analyzed due to weak axonal projections in the target area. **b.** Pooled ODRs from all analyzed coronal sections show an asymmetric distribution around 1. Log ODR values (*inset*), on the other hand, show a more symmetric distribution. **c.** Scatterplots showing the correlation of log ODR values for axonal terminations in all shared target areas of injected pairs of areas. Each point plots against each other the log ODR for the pathway that originates in the pair of injected areas denoted for each graph (at the top and the right of each graph), and which terminates in every common target area. Areas that exhibited weak or absent projections from one of the two injected areas (grey blocks in Fig. 2a) were excluded. A line of unit slope that best fit the points is plotted in each graph. The y-intercept of the unit sloped line is shown in Supplementary Figure 10. Dotted lines, coordinate axes. *Inset*. Log ODR for each pathway plotted against that of its reciprocal counterpart for all 74 pathways that have a dense reciprocal connection (see Fig. 2a). Upper matrix and lower matrix refer respectively to the ODR values in the upper/right and lower/left triangular halves of the matrix in Fig. 2a.

Terminations of projections from areas AM and POR, on the other hand, exhibited the lowest mean ODR values among the ten visual areas indicative of their roles as higher-order association cortical areas. The matrix thus suggests characteristics of a hierarchically organized system of areas in which the tendency to target L2-4 and L1 gradually changes.

Additionally, a hierarchy would be expected to exhibit a systematic and consistent gradient of feedforward and feedback relations between any two of the areas forming the network. More specifically, if a pathway is feedforward in one direction, the reciprocal pathway is expected to be feedback, and likewise the hierarchical separation (i.e. distance) between every pair of areas is expected to be consistent with the hierarchical positioning of all areas across the network. The matrix in Figure 2a shows that out of the 80 pathways, 74 exhibited a reciprocal connection that was dense enough for analysis, i.e. we identified 37 pairs of areas with dense bidirectional connections. To examine if there is any consistent relation between feedforward and feedback pathways, we plotted for all 37 bidirectionally connected areal pairs the log of the ODR in one direction against that in its reciprocal direction (Fig 2c inset). This shows a negative association (*p* = 0.014, F-test; n = 74 pathways) indicating that the more feedforward a pathway is in one direction, the more feedback the reciprocal connection is on average in the opposite direction.

To further investigate the consistency of hierarchical rules across the network, we examined if distance values between any two areas reflected their respective positions within the hierarchy. We reasoned that interareal hierarchical distances would be independent of the injection site, and consistent across injections. For example, in a strict hierarchy, the relative ordering of areas LM and PM, and the hierarchical distance separating them, would remain constant with respect to their laminar termination patterns in each of their target areas, meaning that the ordering is expected to be the same for all shared targets.

Formally, we are looking for a hierarchical distance value *δ* _ij_ between any two areas i and j, which we define as the difference between the respective hierarchical level values (h_i_ and h_j_) of the two areas (Supplementary Figure 10a). Thus,

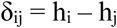

For two cortical areas p and q that are each injected independently, we can estimate the hierarchical distance measure from each of these areas to a common target area i:

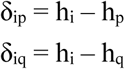

Therefore,

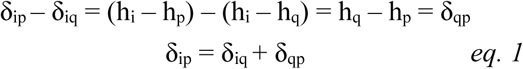

where δ_qp_ is the estimated hierarchical distance between areas q and p. This relation is in the form of a linear function for a line of unit slope in which the intercept gives the hierarchical distance between the two injected areas. Thus, for any target area *i*, a hierarchical distance measure between two injected areas can be estimated by plotting against each other the anatomical indices (such as log ODR) for the respective hierarchical distances between the two injected areas and the shared target area *i*. We therefore plotted log ODR of axonal projections for all pairs of injected areas to their common targets against each other, and fit a line of unit slope to the data to obtain the intercept (Figures 2c, Supplementary Figure 10b). The results indicate that the hierarchical distance between V1 and each of the other areas are, on average, larger than distance values separating any other pair of cortical areas (Supplementary Figure 10b). Area LM similarly appears to be relatively distant from most other areas. The majority of higher areas, however, show relatively low distance values in relation to each other, suggesting they may lie in similarly matched levels, pointing to a lack of distinct, non-overlapping processing stages higher up the visual processing scheme.

### Hierarchy of mouse visual areas

To further probe the hierarchy among higher cortical areas, we used a beta regression model to estimate hierarchical level values for each area (see Methods). The beta distribution, a continuous probability distribution defined on the standard unit interval (0,1), can be obtained from the ODR data through the inverse logistic transformation of the natural log of the ODR values:

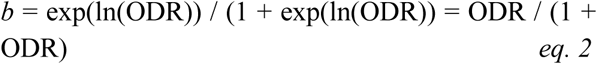

The transformation *b* is thus equal to the proportion of axonal projections in L2-4 to the total axonal projections in L1-4, and has a frequency distribution that lies within the standard unit interval (Figure 3a). The beta regression model was used to estimate the hierarchical level values that best predict ODRs for each cortico-cortical pathway. Area V1 was set at level 0, and the model estimated the values for each area such that the difference between the hierarchical level for any two areas best matched the natural log of the ODR for the connection linking the areas (Figure 2a). The goodness of fit was evaluated by plotting the estimated level values against *b* (Figure 3b, r = 0.87, t(78) = 15.9, p << 0.001; 95% conf. int.: (0.810, 0.917)). Figure 3c shows the estimated hierarchical levels for each area, and with the areas separated into two branches according to the visual processing stream to which they belong^25^. The results reveal that out of the nine higher areas, axonal projections from LM exhibit termination patterns that are most similar to those from V1, reaffirming its role as the second cortical stage for visual processing. Projections from POR and AM had the most dissimilar laminar patterns to those from V1 indicating their role in mediating mainly top-down influences within the visual network.

**Figure 3.**
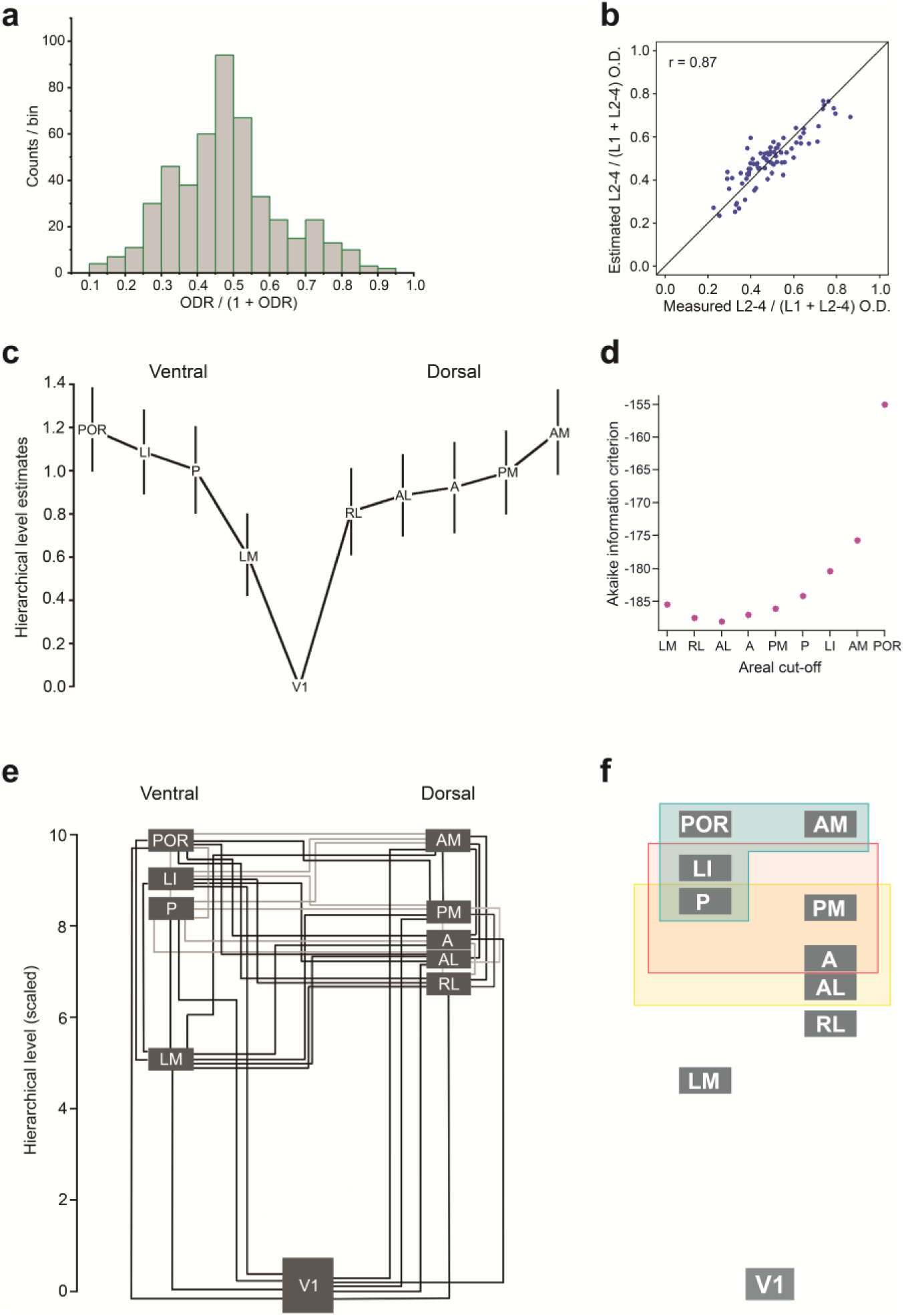
Construction of mouse visual cortical hierarchy. **a.** Distribution of the inverse logit transformed values of ODRs for all pathways (see eq. 2) **b.** Hierarchical distance values for all pairs of areas estimated by a beta regression model show a high goodness of fit with the measured values (r^2^ = 0.76). **c.** Estimated hierarchical levels obtained by a beta regression model such that the level value of V1 is set at 0, and differences between any two hierarchical level values best predict the ODR for the pathways connecting the respective areas. Vertical lines demarcate 90% confidence intervals. The areas have been divided into previously described dorsal and ventral streams^.25,27^ **d.** The AIC values for nine models in which the highest *n* areas (*n* going from 1 to 9, beginning with the highest area POR) were constrained to be part of the same level, and the beta regression fit performed for each such model. The lowest AIC value occurs for the model in which all areas from AL and higher are combined into a single level, indicating that the model in which V1, LM, RL, and a fourth level containing all other areas are treated as four different levels, is the hierarchical model with the best predictive power. **e.** Hierarchical levels similar to Figure 3c, but scaled to values between 1 and 10. Black lines interconnect pairs of areas that show a significant difference in their hierarchical levels. Grey lines interconnect areal pairs that lack a statistical significance in their hierarchical level. **f.** Illustration of the overlapping hierarchy of the network. All pairs of areas within each box lack a statistically significant hierarchical separation, and pathways interconnecting these areas can therefore be considered to be lateral (i.e. neither feedforward nor feedback).

How many different levels compose the hierarchy? To identify the number of distinct stages that best describes the processing sequence, a cumulative procedure was employed in which the highest *n* areas (*n* going from 1 to 9), when arranged in the increasing order of their hierarchical levels from V1 to POR (Figure 3c), were constrained to be part of the same level. This was done by adding the next lowest column to the *n* highest columns, starting with POR as the highest area and performing the beta regression fit for each such model. This generates a set of nested models. The best model was assessed as the one with the lowest Akaike Information Criterion (AIC) value^24,31^, which identifies the model with the best predictive power for the inclusion of new data. This was the case in which all areas from AL and above were treated as being part of the same level (Figure 3d).

Similarly, the AIC was lower for the model in which areas LM and RL were treated as separate levels than when they were combined into a single level. The model in which V1 was combined with LM gave a much higher AIC. Together, these results point to an organization of four levels – V1, LM, RL, and all other areas combined – as the best model describing the processing pathway.

Level values were estimated for each area relative to each of the other nine areas to evaluate whether the difference between the hierarchical levels of any two areas reflected a statistically significant separation. The organization of the areas based on such pairwise examinations is displayed in Figure 3e in which black lines connect areas that are at significantly distant levels (*p* < 0.05, Tukey’s range test), and grey lines connect areas whose estimated level values lacked a statistically significant difference; the latter can accordingly be considered to indicate lateral connections. This shows that V1 and LM are significantly separated from all other areas based on the beta regression model, while area RL did not show a significant separation from areas AL and A (Figure 3e). Higher areas were characterized by a large number of inconsistencies in their hierarchical ordering. For example, while the analyses indicated area PM as being lower than areas AM and POR and residing at an equal level with area P, P did not exhibit projection patterns that were significantly different than those projecting from AM and POR (Figure 3e). Additionally, multiple higher-order areas could be grouped such that every areal pair within the group lacked a significant difference in their hierarchical positioning (Figure 3f). The hierarchy can therefore be thought of as comprising overlapping processing levels at higher stages, each characterized exclusively by lateral connections. In contrast to these higher-order areas, all connections emerging from or terminating in V1 and LM can be classified as being either feedforward or feedback pointing to a conventional hierarchical organization in early visual representation. Importantly, despite the overlapping characteristic of the overall network, there are distinct processing routes from V1 to POR or AM, each comprising areas at significantly separated levels (Figure 3e). The findings thus indicate the presence of multiple, canonical systems embedded within noncanonically organized networks.

### Receptive field size across the visual cortical hierarchy

Because areas at increasingly higher levels may be expected to integrate inputs from an increasingly larger number of lower areas, an expected functional consequence of a cortical hierarchy is that neurons in higher areas display larger receptive fields than those in lower areas. To test if the hierarchy established on anatomical rules is consistent with the proposed summation of visual space within such a system, receptive fields were mapped for neurons in each of the ten areas in response to drifting gratings in anesthetized mice (n = 230 neurons from 23 mice; Figure 4). The drifting grating stimulus (5 deg circular patch) was flashed at multiple locations of a display during recordings of single unit responses. Representative response fields are shown in Figure 4a. To analyze the size of receptive fields (RFs), the recorded response fields were fit with a two-dimensional Gaussian. RFs were measured as the contour corresponding to ±2 SDs of the fitted Gaussian, and the RF size was calculated as the diameter of a circle with the same area as that of the ellipse formed by the major and minor axes of the Gaussian.

**Figure 4.**
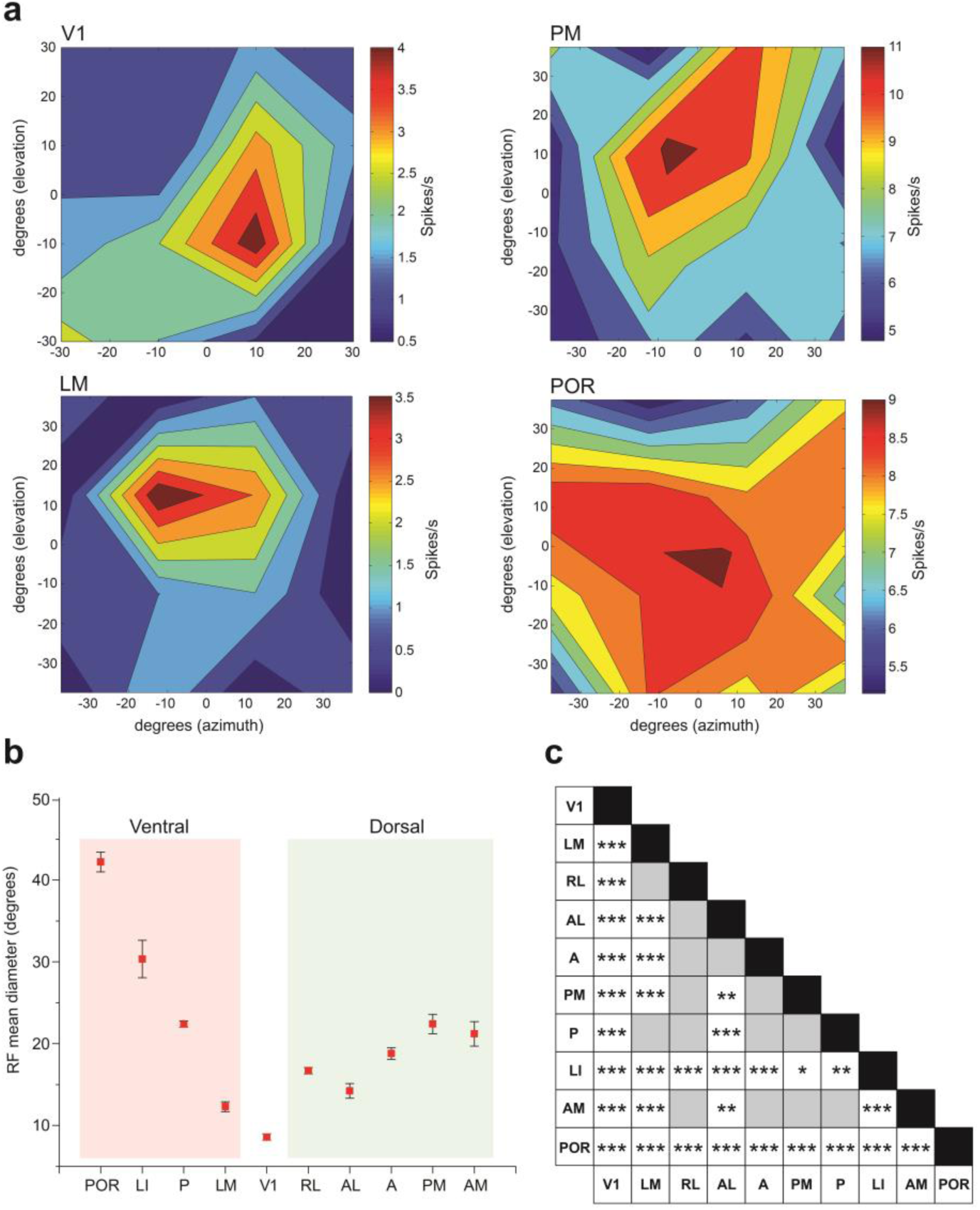
Receptive field diameters scale with hierarchical levels within each processing stream. **a.** Representative heat maps of firing rates of neurons in V1, LM, PM, and POR, used to determine RF diameters. **b.** Within each processing stream, RF diameters show an overall increase in areas at increasingly higher hierarchical levels (*p* < 0.001 for both dorsal and ventral streams, one-way ANOVA). This effect is more prominent in the ventral stream. **c.** Statistical significance of RF diameters for all pairs of areas. \**p* < 0.05, \*\**p* < 0.01, \*\*\**p* < 0.001. Tukey’s range test. Grey blocks indicate no statistical significance.

Individual visual areas have been previously demonstrated to preferentially form reciprocal connections with higher projection weights with areas belonging to either a dorsal or a ventral network^25^, indicating that the cortical hierarchy involves at least two streams of preferentially interconnected areas. We therefore separated the nine higher areas into the two streams to examine if the RF diameters showed a successive increase within each branch of the hierarchy. Mean RF diameters depended on both hierarchical organization as well as on processing stream (Figure 4b). While both dorsal and ventral streams exhibited overall increases in RF diameters with increasing hierarchical positioning (*p* < 0.001 for dorsal stream, *p* < 0.001 for ventral stream, one-way ANOVA), this relationship was more pronounced in the ventral stream (Figure 4b). Ventral stream areas LI and POR had the largest RF diameters (30.35 ± 2.3 degrees, n = 9, and 42.2 ± 1.2 degrees, n = 6, respectively) out of all the areas, significantly larger (Tukey’s range test, Figure 4c) than the RFs of each of the two highest dorsal stream areas AM (21.2 ± 1.5 degrees, n = 9) and PM (22.4 ± 1.2 degrees, n = 6). Areas V1 (8.59 ± 0.4, n = 86) and LM (12.3 ± 0.6, n = 72) had the smallest mean RF diameters (Figure 4c), consistent with their respective positions at the bottom of the visual cortical hierarchy. Together, these results indicate that in each processing stream hierarchical positioning determined by anatomical criteria is largely consistent with spatial summation, pointing to a key role of higher-order areas in integrating spatiotemporal information emanating from lower areas.

## DISCUSSION

Cortical hierarchies determined by structural consistencies provide insight into how feedforward, feedback, and lateral connections assemble incoming afferent inputs into complex representations and how they update the internal model of the visual scene^8,33,34^. Top-down influences are thought to underlie various functions of the visual system including feature binding, scene segmentation, object-based attentional selection, working memory, and decision making^35,36^, implying a critical role of a hierarchical arrangement of areas in visual perception. Genetically and physiologically tractable mammalian species such as the mouse are particularly useful for identifying fine-scale, cell type-specific circuit motifs that vary across functionally specialized interareal pathways^14,21,37^. The mouse visual connectome, however, is ultra-densely organized: 97% of all possible combinations of any two cortical areas have been estimated to be connected, and within the ten visual areas described in this study, the density rises to ∼99%^24^. Such a high density raises the question of whether functionally diverse areas, each with distinct spatiotemporal response properties, are ordered such that there exists a sequence of higher-order areas that each integrates inputs from an increasing number of lower areas. Our analyses of the laminar termination patterns of anterogradely labeled axonal projections within the network of these ten areas indicate that while a hierarchy emerges from the differential laminar distribution of projection weights, the network involves a system that can be described as an overlapping hierarchy comprising four processing levels: V1, LM, RL, and the remaining seven areas forming a fourth processing stage in which canonical circuits are embedded within a noncanonical network. For example, areas AL, A, PM, and P form a laterally interconnected network that is lower than AM and POR. We suggest that such an organization could underlie a processing strategy in which a shared property of the four lower areas, such as a motion-based spatial reference frame, may be differentially employed by AM and POR for goal-directed behavior.

Studies in primate cortex indicate that signals emerging from V1 are transmitted to higher areas through broadly two processing streams, each thought to underlie distinct visual functions^38,39^. The dorsal stream, which preferentially interconnects V1 with areas in the posterior parietal lobe, is thought to be responsible for spatial navigation and the visual guidance of action. The ventral stream, on the other hand, which routs output signals from V1 to the temporal lobe, is considered to mediate object recognition^38,40^. Embedded within the densely interconnected network of mouse visual areas are two subnetworks each comprising areas that preferentially interconnect with each other, leading to the proposal that the mouse visual system, much like its primate analog, also utilizes distinct processing streams for visual function^25^. Here we show that the ventral stream area LM and dorsal stream area RL are low in the hierarchy, with POR and AM at the highest levels of each stream. We find that projection strength does not predict hierarchical level: AL has a stronger input from V1 than does RL, yet AL is at a higher level than RL. While both AL and RL are specialized for processing visual motion^41^, the two areas appear to underlie distinct functions; AL was recently shown to more strongly respond to coherent motion within random dot patterns^42^, while RL has been shown to be specialized for processing global pattern motion^43^, visuotactile multisensory integration^44^, and for the routing of information from a subpopulation of direction-selective retinal cells^45^. RL also forms overall stronger connections than AL with somatosensory and motor areas^24^. Together, these results suggest the existence of hierarchically organized substreams within the dorsal stream, analogous to that proposed in primate cortex^40^: AL being the gateway of a parieto-temporal pathway, and RL linked more strongly with a parieto-premotor/prefrontal pathway.

Interestingly, across streams, relative RF sizes were not indicative of hierarchical levels; RF diameters were consistent with hierarchical levels only within a stream. For example, despite AM and POR being at similar levels, each at the top of its respective stream, the RF diameter of POR was significantly larger than that of AM. The ventral stream area LI also exhibited a larger RF diameter than AM despite being hierarchically lower (Figure 4b, c), reminiscent of primate cortex where ventral stream areas have larger RF diameters than their dorsal stream counterparts^46,47^. The overall larger RF sizes in the ventral stream may reflect a crucial specialized function of the subnetwork of ventral stream areas in integrating information from a larger spatial extent of the visual field for the processing of motion detection and for mapping the local space in the allocentric frame of reference^48,49^. Notably, this is different from the role the ventral stream is believed to play in the monkey where it is specialized for the processing of textures and shapes^38^.

The hierarchical organization described in this study has dissimilarities with the recently published network study that expanded the Allen Mouse Brain Connectivity Atlas^14^. While both studies showed V1 at the bottom, and areas AM and POR at the top of the hierarchy of visual areas, there are differences in the positioning of other areas perhaps most notably that of LM and RL as the second highest level of the processing pathway. Such disparities are attributable to projections to areas outside of the ten visual areas examined in this study, as well as in differences in the analysis used to construct the hierarchy and the exclusion of projections from areally overlapping injection sites^24,50^. Importantly, both studies conclude that the hierarchy of the mouse visual cortex is shallow, with few distinct processing levels. The shallowness of the hierarchy likely reflects the larger RF sizes in mouse V1, compared with that of primates, and that it takes only four instead of ten steps up the hierarchy for pooling visual space into a 40 deg RF, as seen in the medial superior temporal area near the top of the cortical hierarchy in macaques^46^.

The ODR metric used here avoids pigeonholing individual projections into feedforward or feedback categories, and more directly estimates hierarchical distance values through a mathematical transformation of the ODR and regression fitting. This avoids violations of feedforward/feedback relationships between a reciprocally connected pair of areas - for example, having such a pair exhibit feedforward or feedback termination patterns in both directions - and allowed us to tease apart the hierarchical characteristics of the network of higher areas. However, it is notable that in numerous cases, projections in both directions of a reciprocally connected pair strongly targeted L2-4, which contain the cell bodies of pyramidal cells; such projections can presumably strongly activate postsynaptic neurons to fire. The existence of such pairs of areas appears to contradict the “no-strong-loops hypothesis”, which proposes that looped interareal circuits comprising only strong, ‘driving’ connections do not exist as they would result in a positive feedback loop resulting in uncontrolled excitation across areas^51^. Possible mechanisms to avoid runaway, epileptic excitation within such ‘strong loops’ involve the differential engagement of inhibition by different pathways^21,37,52^, as well as through dynamically changing, differential levels of input strengths across layers depending on context and visual salience. The latter mechanism can be formalized in the posterior parietal cortex^24^ as a coordinate transformation^53^, a process in which one functionally specialized coordinate frame is transformed to another depending on the sensorimotor function that needs to be achieved^54^. In a predictive processing framework, the observations described in this study implicate each higher-order area in providing predictive signals to multiple other areas, potentially underlying parallel processing pathways for the representation of diverse features of a scene. Consistent with this suggestion, it was recently proposed that a strict hierarchy of areas is not required for predictive processing of sensory signals - a pair of areas can conceivably exchange predictions and prediction errors in both directions, depending on the sensory modality in which predictions and prediction errors are being generated^54^. We extend this proposition to the representation of different features involved in visual function and spatial navigation, and suggest that the organization of laterally interconnected areas with different response properties does not imply a lack of canonically hierarchical processing, or the presence of functional ‘strong loops’, but that the functional hierarchy required for achieving a specific visual function dynamically changes as a function of context and the salience of visual inputs. In other words, maintaining and updating internal representations of complex features higher up the hierarchy may require coordinate transformations that selectively recruit one particular functional hierarchy out of many possibilities within the noncanonical network.

While the focus of this study was primarily the intracortical network, the essential role of the thalamus in regulating signal flow across visual cortex must be considered in the description of cortical function. The pulvinar, a higher-order thalamic nucleus, is crucially involved in controlling feedforward and feedback communication within the visual interareal hierarchy^.55,56^ In the mouse, the pulvinar is also likely a critical component in the processing of retinal signals by the higher-order area POR. POR was recently shown to be primarily activated not through signals emerging from V1, but rather almost exclusively from signals transmitted through the collicular-pulvino-cortical pathway^57^. This indicates that V1 is not the exclusive gateway into the cortical hierarchy, but that visual signals can enter the cortical network at higher processing stages through the pulvinar. Additionally, anatomical tracing and MAPseq experiments suggest that the routing of output signals from V1 is constrained by preferential targeting of subsets of areas by axon collaterals of individual projection neurons residing in V1^58^, which may imply the broadcasting of correlated signals from individual V1 neurons to multiple levels of a processing hierarchy, as well as to areas belonging to different processing streams. Thus, deducing the algorithms underlying cortical processing requires a thorough understanding of thalamic connectivity with distinct cortical areas, as well as of the most efficient direct and indirect paths between areas for signal transmission across the hierarchy^25^.

## Supporting information

All Supplementary figures

## SUPPLEMENTARY FIGURE LEGENDS

**Supplementary Figure 1.**

a. Rostrocaudal series of coronal sections of left hemisphere in which AL (denoted by asterisk) was injected with BDA. Darkfield images of anterogradely labeled axonal projections (yellow/orange) to distinct visual areas (white squares). Numbers denote the coronal plane corresponding to the respective rostrocaudal location shown in the inset. Fluorescent images of retrogradely labeled callosal neurons (light cyan), after injection of bisbenzimide into the right hemisphere, aid in the identification of areal borders. *Inset, In situ* image of left hemisphere, before coronal sectioning, showing retrogradely labeled callosally projecting neurons (light cyan). Asterisk denotes injection site in AL. Horizontal lines and numbers denote the coronal planes shown above. Scale bars, 1 mm.

b. High magnification images of regions within the white squares in Supplementary Figure 1a. Axonal projections from AL target all nine areas with varying strengths, and are observed in all six layers. Scale bar, 200 µm.

**Supplementary Figure 2**

Darkfield images of anterograde BDA labeled axonal projections from LM to areas V1, AL, RL, PM, P, A, LI, AM, and POR. LM axons preferentially target L1 of V1, and show varying preferences in the other areas, often terminating densely in L2-4 in higher areas. Asterisk in the LI panel denotes region in LM near injection site.

**Supplementary Figure 3**

Darkfield images of BDA labeled axonal projections from PM to different layers of V1, LM, AL, P, A, LI, AM, and POR. Note that PM projections to RL were too weak for analysis. Arrowheads in the panels for areas A and LI demarcate respective boundaries used for analysis. The respective locations of AM and LM are shown in the A and LI panels. Scale bar, 200 µm.

**Supplementary Figure 4**

Darkfield images of BDA labeled axonal projections from RL to different layers of V1, LM, AL, PM, A, LI, AM, and POR. Scale bar, 200 µm. Arrowheads demarcate borders of POR.

**Supplementary Figure 5**

Darkfield images of BDA labeled axonal projections from P to different layers of V1, LM, AL, PM, A, LI, and POR. Arrowheads demarcate area LI. Locations of LM and P are shown in the LI and POR panels, respectively. Scale bar, 200 µm.

**Supplementary Figure 6**

Darkfield images of BDA labeled axonal projections from LI to V1, LM, AL, RL, PM, P, AM, and POR. Arrowheads demarcate borders of areas LM, RL and POR used for analysis. LI is depicted in the LM and POR panels. Scale bar, 200 µm.

**Supplementary Figure 7**

Darkfield images of BDA labeled projections from A to V1, LM, AL, RL, PM, LI, AM, and POR. Arrowheads demarcate borders of AM. Position of A is shown in the AM panel. Scale bar, 200 µm.

**Supplementary Figure 8**

Darkfield images of BDA labeled axonal projections from AM terminating in different layers of areas V1, LM, AL, RL, PM, P, LI, A, and POR. Arrowheads demarcate borders of RL and LI used for analysis. Projections to AL and LM are shown in the RL and LI panels, respectively. Scale bar, 200 µm.

**Supplementary Figure 9**

Darkfield images of BDA labeled projections from POR to areas V1, LM, AL, RL, PM, P, LI, A, and AM.

**Supplementary Figure 10**

a. Illustration of the relationship between hierarchical levels (h) and distances (*δ*), and the derivation of eq. 1.

b. Each block shows the y-intercept of the respective graph of Figure 2c (triangular plot). Each value acts as a measure of hierarchical distance (see eq. 1) between the two areas denoted at the top and the right.

## ACKNOWLEDGMENTS

This work was supported by the National Eye Institute (R01 EY016184, R01 EY020523, R01 EY022090, and R01 EY027383 to A.B.), and LABEX CORTEX (ANR-11-LABX-0042 to H.K.) of Université de Lyon (ANR-11-IDEX-0007) operated by the French National Research Agency (ANR), (ANR-15-CE32-0016 CORNET to H.K.), (ANR-17-NEUC-0004 to H.K.), (A2P2MC to H.K.), (ANR-17-HBPR-0003, CORTICITY to HK), (ANR-19-CE37-0000, DUAL_STREAMS to K.K.).

## AUTHOR CONTRIBUTIONS

Q.W. and A.B. designed experiments; Q.W., R.D.D., W.J. and A.B. performed experiments; R.D.D., K.K., A.M., and W.J. analyzed data; R.D.D., A.B., K.K., and H.K. wrote the manuscript;

