## Supplementary figures and images for "Canonical and noncanonical features of the mouse visual cortical hierarchy"

### All Supplementary figures

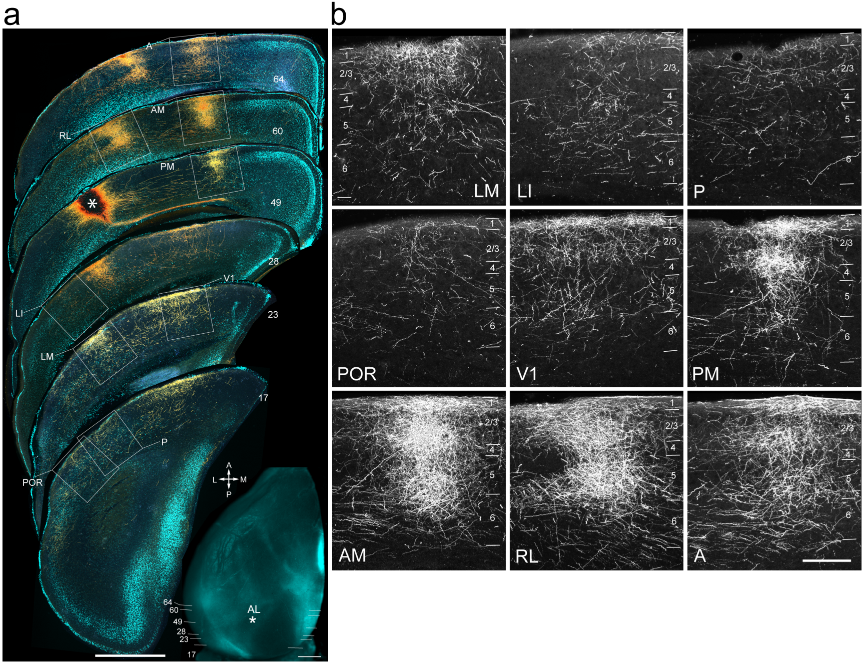

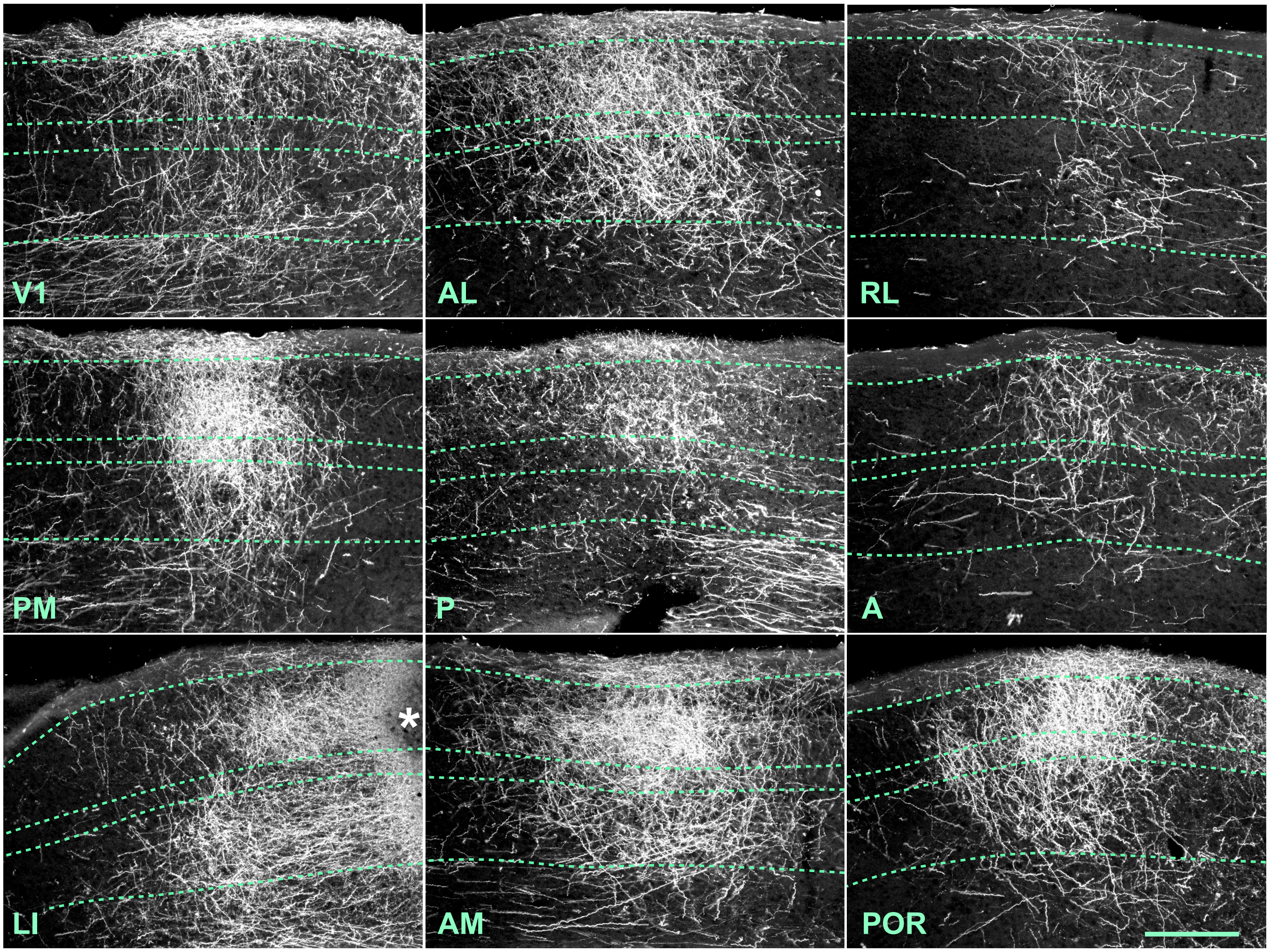

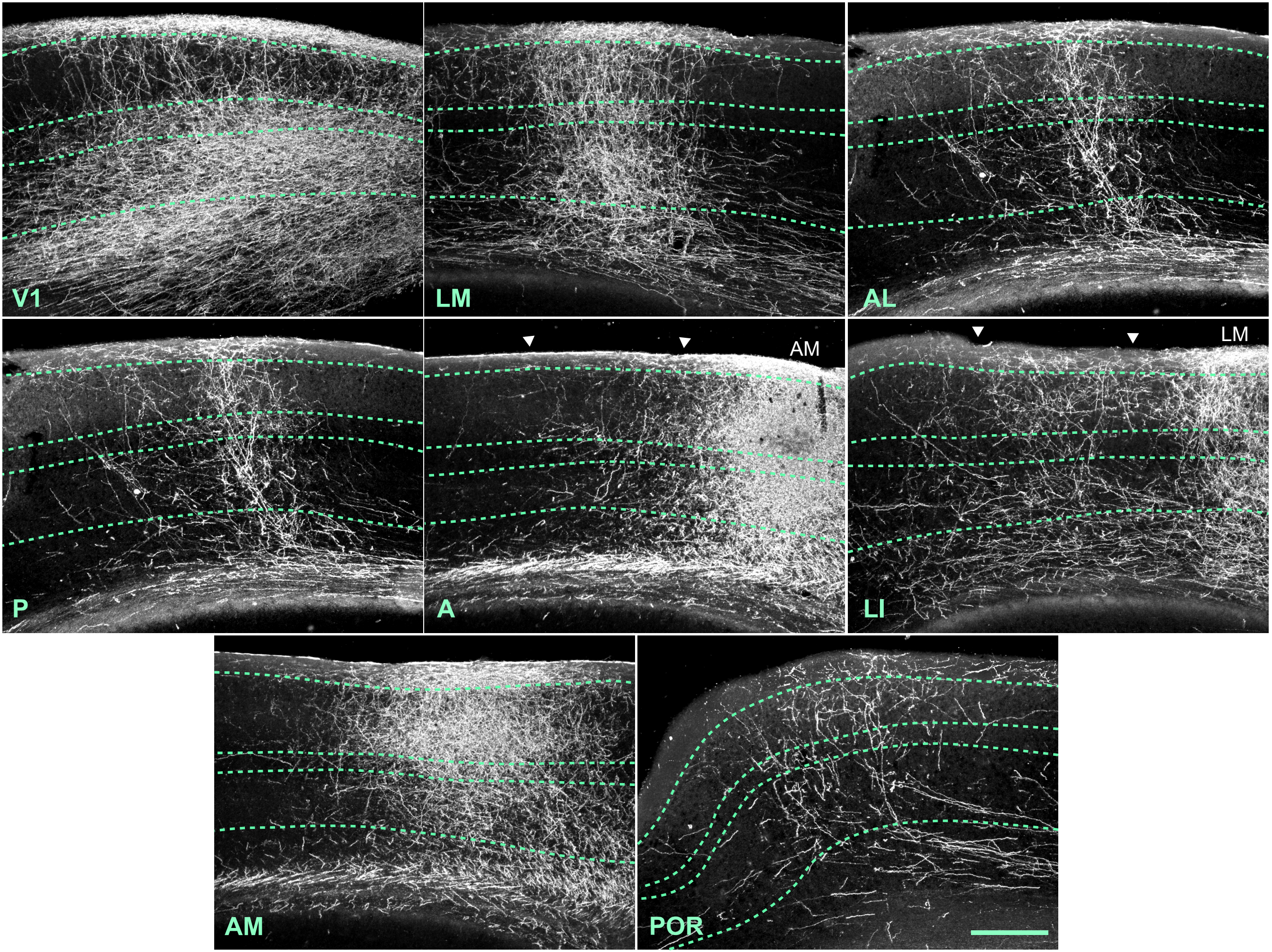

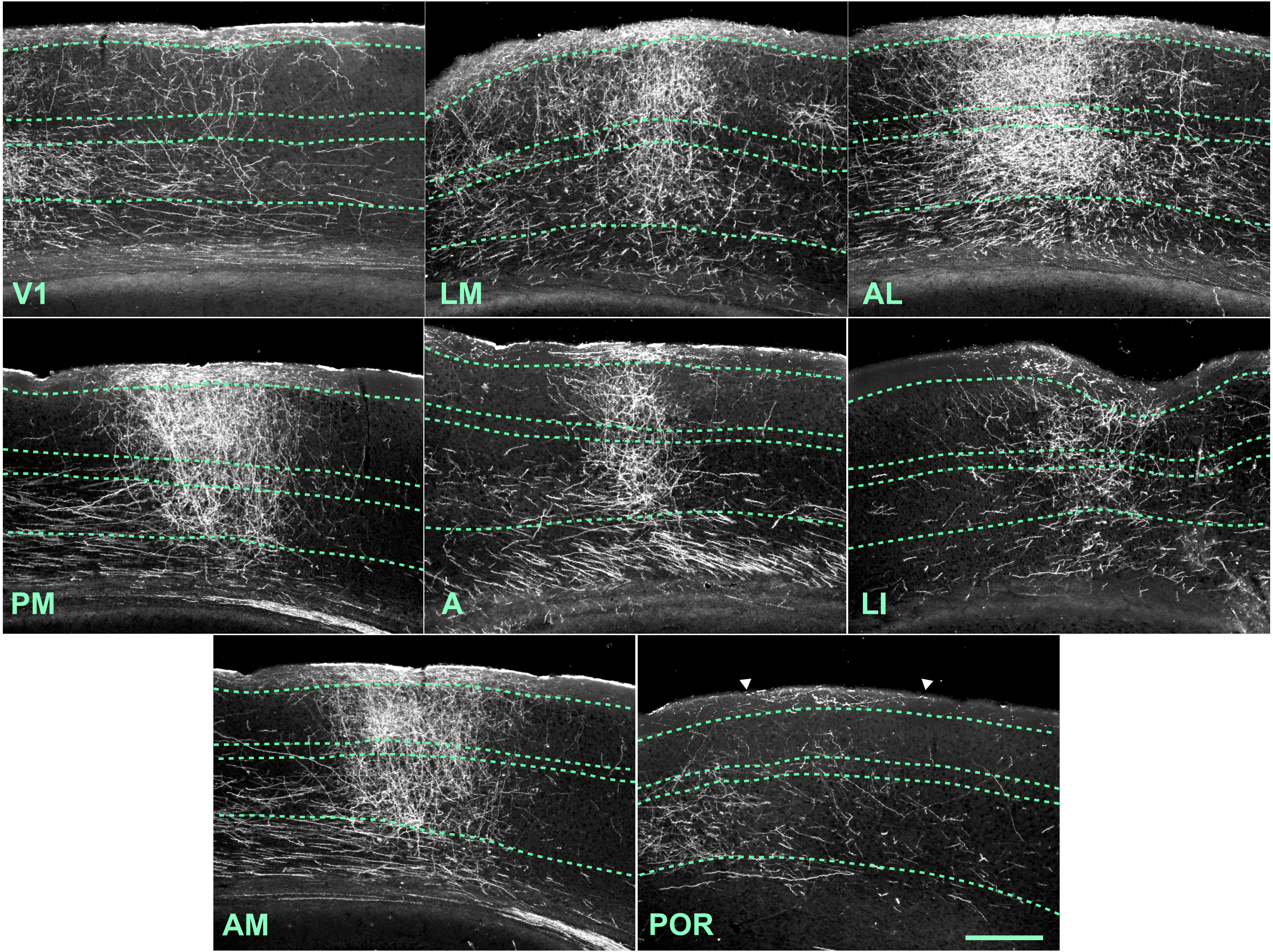

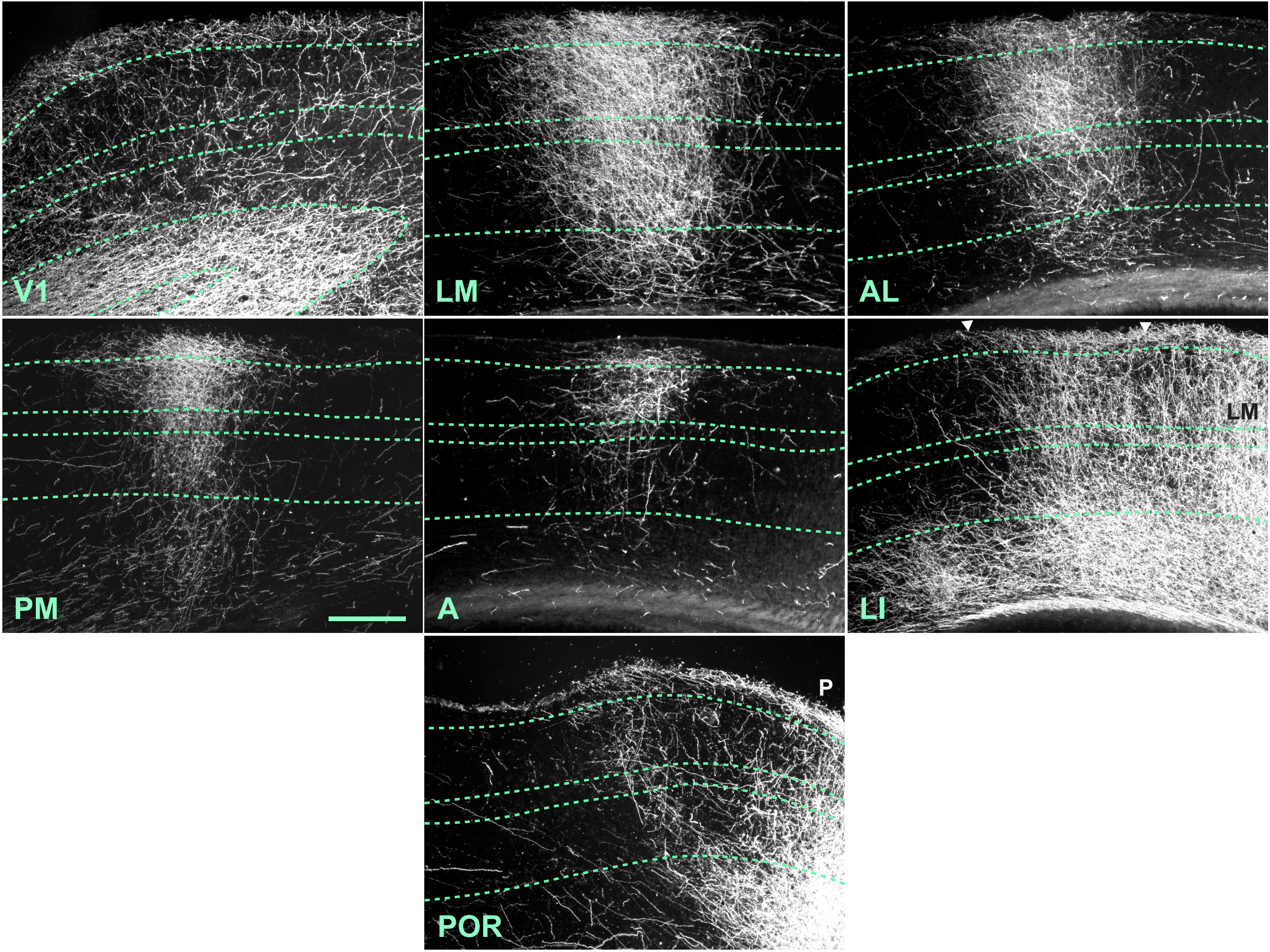

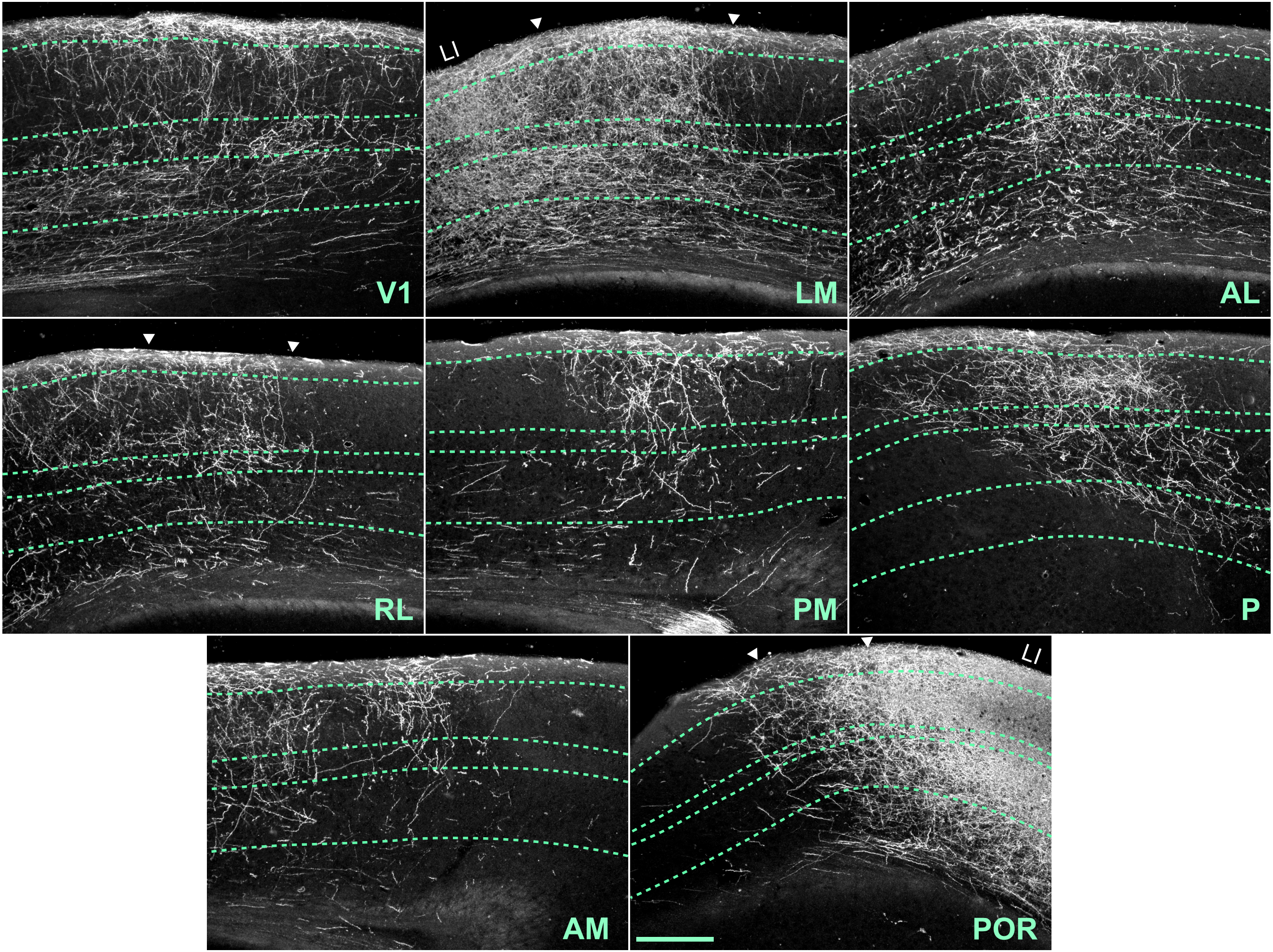

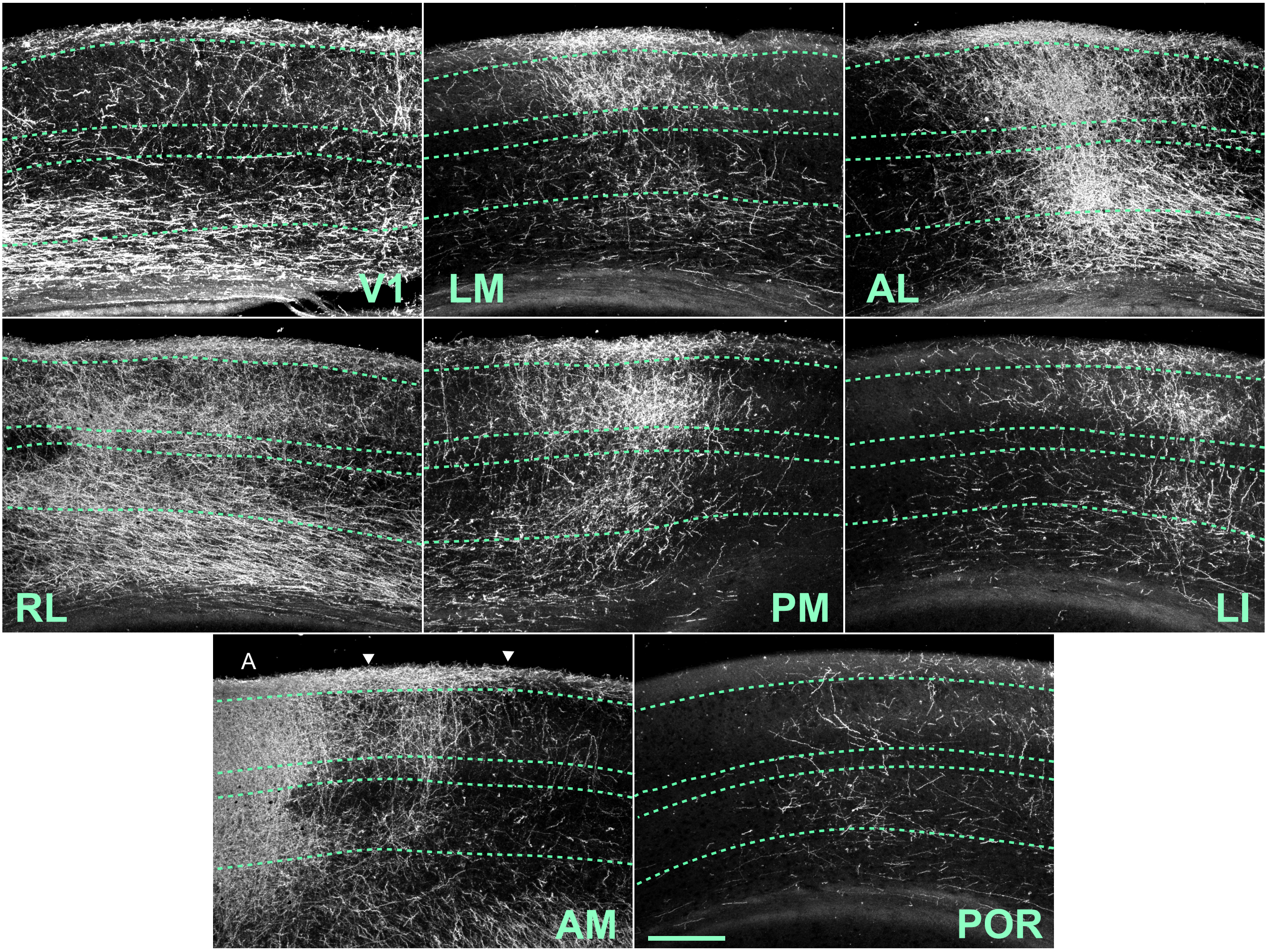

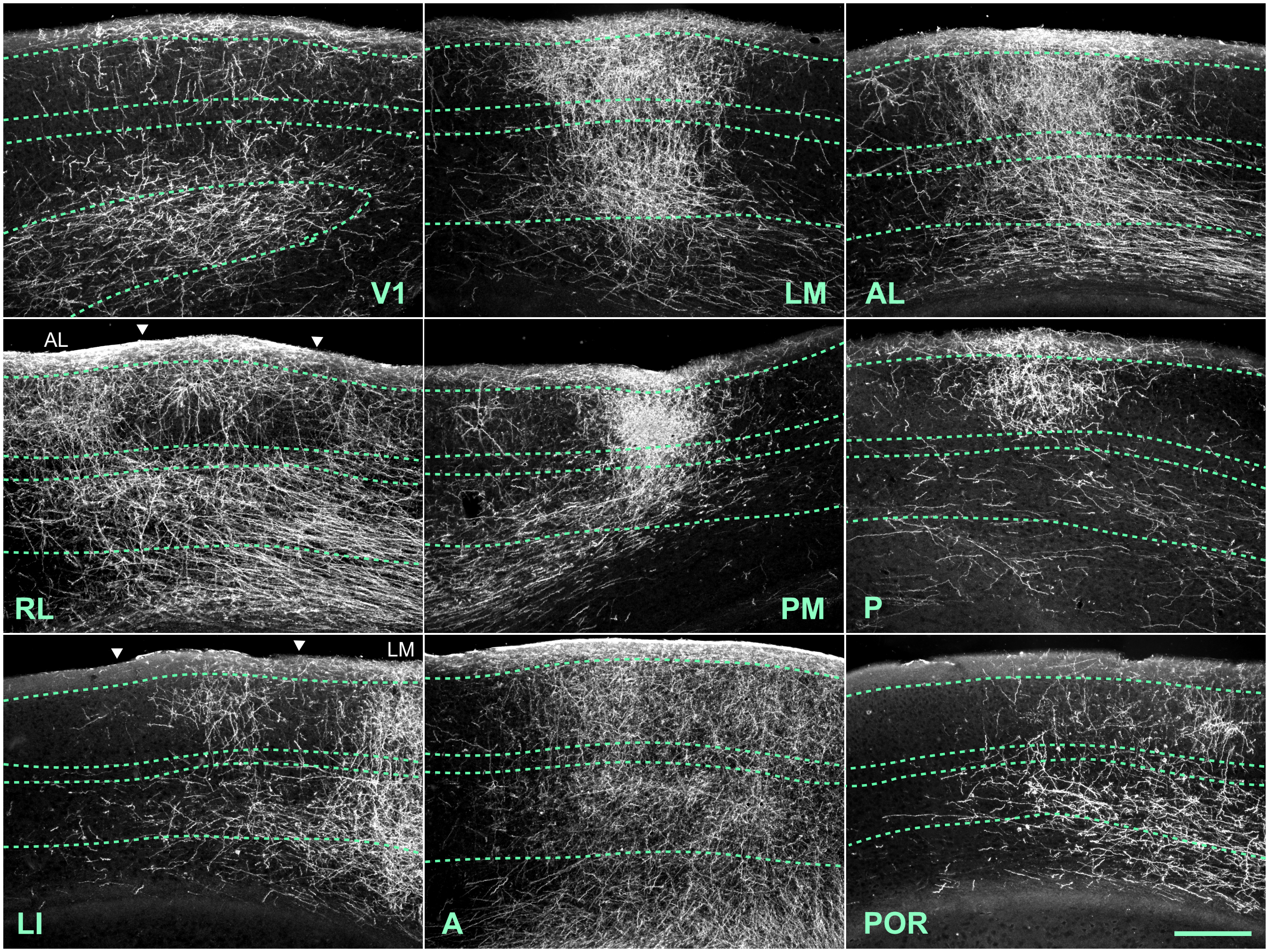

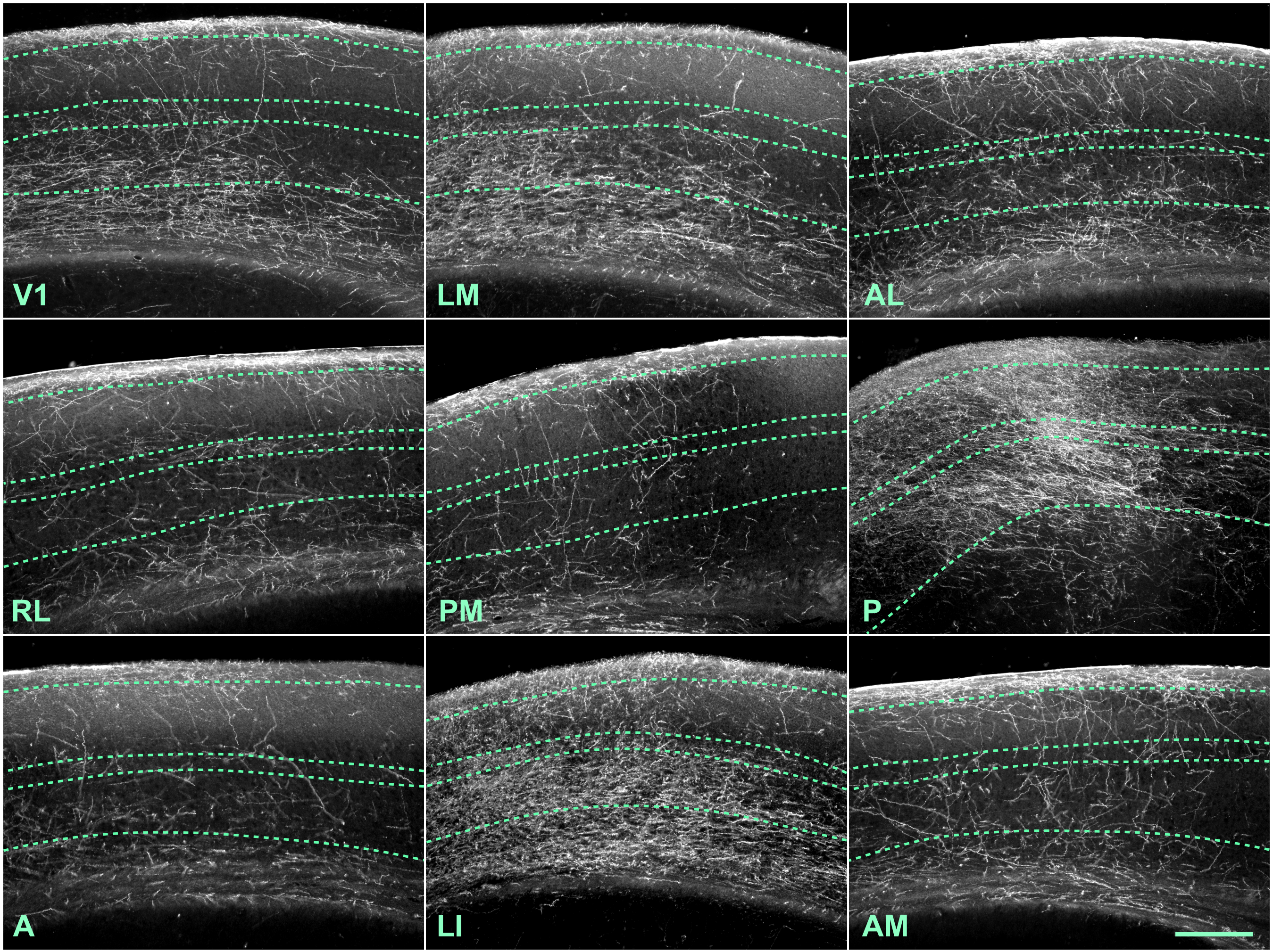

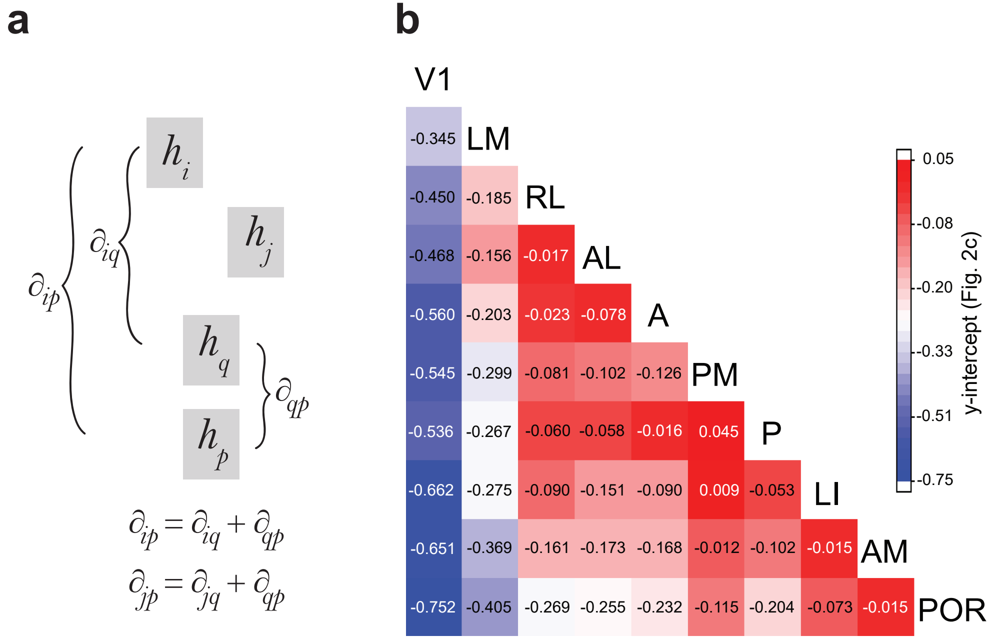
